# Metabolic ecology and habitat stability explain the disproportionately high species richness in standing waters

**DOI:** 10.1101/2025.10.23.683910

**Authors:** Laura Anna Mähn, Christian Hof, Seth Bybee, Roland Brandl, Stefan Pinkert

**Affiliations:** Department of Animal Ecology, Phillipps-Universität Marburg, Marburg, Hessia, Germany; Department of Global Change Ecology, University of Würzburg, Würzburg, Bavaria, Germany; Department of Biology and Monte L. Bean Museum, Brigham Young University, Provo, Utah, USA; Department of Conservation Ecology, Phillipps-Universität Marburg, Marburg, Hessia, Germany

**Keywords:** Body size, climate stability, dragonflies and damselflies, environmental drivers, freshwater organisms, habitat-stability-dispersal hypothesis, lentic and lotic habitats, Metabolic Theory of Ecology, range size, species richness

## Abstract

The Metabolic Theory of Ecology (MTE) conceptualizes that temperature is the primary driver of species richness, a pattern well supported in terrestrial taxa but less certain for freshwater organisms. Limited global-scale evidence and frequent violations of MTE’s assumptions, particularly the stationarity of body size and abundance, further obscure its applicability. In freshwater systems, body size and abundance are tightly linked to dispersal and range size, which differ markedly between running-water (lotic) and standing-water (lentic) species, as proposed by the Habitat-Stability–Dispersal Hypothesis (HSDH). Adaptations to habitat stability may therefore generate distinct biogeographical trait patterns and modify richness–temperature relationships predicted by MTE. Utilizing comprehensive global functional, phylogenetic, and distributional data on dragonfly and damselfly species (83%) and habitat information (46%), we tested MTE predictions for lentic versus lotic species. Lotic species richness followed MTE expectations (slope: –0.469) more closely than lentic species richness (slope: –0.283). The proportion of lentic species in an assemblage was the strongest predictor of deviation in the species richness-temperature relationship (R^2^ = 38%). Assemblages dominated by lentic species clustered in climatically unstable regions and mainly including smaller-bodied species with larger ranges. Phylogenetic comparative analysis shows a strong phylogenetic signal in habitat preference, with the most species rich and northernly distributed families comprising predominately lentic species. Our findings suggest that adaptations to habitat stability facilitated the colonization and persistence of lentic species in harsh and fluctuating climates both past and present causing largely divergent species richness patterns of lentic and lotic odonates. Integrating HSDH-related traits (body and range size) not only substantially improves the explanatory power of the MTE, but also reveals a trait syndrome with broad implications for the biogeography and climate change responses of freshwater communities.

## INTRODUCTION

Understanding the drivers of species richness is pivotal for anticipating species’ responses to climate change and effective biodiversity conservation (Gaston 2000, Mittelbach et al. 2007, Rahbek et al. 2019). Strong relationships between species richness and temperature, water availability as well as productivity, documented for a broad array of regions and organisms suggest the existence of general mechanisms driving species’ distributions and diversity patterns (Hawkins et al. 2003, Currie et al. 2004, Rahbek et al. 2019). Many of the proposed mechanistic explanations address fundamental physiological principles of resource availability and species’ energy demand (Hawkins et al. 2003, Currie et al. 2004, Algar et al. 2007). The Metabolic Theory of Ecology (MTE) is a promising framework for species richness conceptualizing an ubiquitous, positive temperature-species richness relationship as the consequence of metabolic energy demands and resource availability (Brown et al. 2004, Algar et al. 2007). Moreover, the MTE allows to derive an expected slope and intercept of this relationship, providing a quantitative baseline to assess how natural selection acting on physiological requirements shapes species richness among organisms and spatial scales (Allen et al. 2002, Brown et al. 2004). The slope is theoretically expected to fall between – 0.60 and –0.70, reflecting how strongly diversity scales with temperature, while the intercept can be interpreted as a measure of baseline diversity at a given temperature, influenced by ecological and evolutionary processes beyond metabolism (Allen et al. 2002, Brown et al. 2004).

Support for the MTE is largely limited to regional and continent-wide studies of terrestrial endothermic animals and plants (Algar et al. 2007, Hawkins et al. 2007, Bailly et al. 2014). Challenges to the applicability of the MTE seem to prevail in isolated habitats, such as mountains, that stand out as centers of diversity (Rahbek et al. 2019), particularly for insects and other ectothermic animals (Buckley et al. 2012, Pinkert et al. 2020b, Pinkert et al. 2025). For example, some of the highest species richness values relative to temperature and productivity cluster in mountain regions. These hotspots challenge the predictions of the MTE, with species richness being higher than expected based on temperature alone (Rahbek et al. 2019, Pinkert et al. 2025a). Notably, mountains and other isolated habitats such as islands often harbour high concentrations of endemic species characterized by low dispersal ability and small range sizes. This pattern suggests that dispersal-diversification dynamics act as major additional constraints on species richness (Brown et al. 2004, Rahbek et al. 2019, Pinkert et al. 2020b, Pinkert et al. 2025a).

Like mountain ranges in a matrix of lowlands or islands in the sea, freshwaters and particularly standing waters are highly isolated habitats with dispersal-diversification dynamics being one of the leading causes of the outstanding species richness they support (Dijkstra et al. 2014). Freshwater ecosystems cover ∼1% of Earth’s surface, yet they host at least 10% of all described animal species, of which 60% comprise aquatic insects (Balian et al. 2008). The two main types of freshwater systems, lentic (standing) and lotic (running) waters, are characterized by distinct differences in their spatial and temporal stability (Ribera and Vogler 2000, Arribas et al. 2012). Lentic water bodies (e.g., ditches, phytotelmas, ponds and lakes) are generally more ephemeral and spatially isolated, with a high probability of drying up seasonally or disappearing over decades to centuries (Bohle 1995, Marsh and Fairbridge 1999). In contrast rivers and streams (lotic waters) often maintain their course and flow regime for thousands to millions of years, making them highly persistent in time and space. The habitat-stability-dispersal hypothesis (HSDH) proposes that the temporal and spatial instability of habitats shapes the evolution of species’ dispersal-related traits (Southwood 1977). In stable habitats, species tend to invest less in dispersal, as local persistence is favored. In contrast, in instable habitats, species evolved traits that enhance persistence, dispersal and recolonization in response to the transient habitat conditions (Southwood 1977, Bohle 1995, Marsh and Fairbridge 1999, Ribera and Vogler 2000, Arribas et al. 2012, Hof et al. 2012, Grewe et al. 2013).

Here, habitat stability refers not to short-term physical variability (e.g. turbulence or changes in flow), but to the long-term seasonality and paleoclimatic instability that require strong dispersal for the colonization of new habitats. Although the HSDH is supported by several comparative studies across freshwater insects, its generality remains debated. Critics argue that dispersal traits may evolve under multiple selective pressures unrelated to habitat stability (e.g. predation) and that empirical tests often struggle to disentangle habitat stability from correlated ecological gradients (Hof et al. 2012, Lancaster & Downes 2018).

In the context of long-term climate stability, i.e. climatic fluctuations over millennial timescales, especially during the Pleistocene, represents a critical filter, with the distributions of species with lower dispersal ability still reflecting past climatic regimes more than adaptations to present day conditions (Pinkert et al. 2018; Pinkert et al. 2025b). These dynamics are particularly obvious but not limited to freshwater ecosystems, where the necessity for frequently (re)colonization of lentic habitats selected for species with greater dispersal ability and ecological tolerance. Indeed, regional-scale support for odonates, caddisflies, and water beetles of North America and Europe suggests that lentic species have larger ranges and show more rapid range shifts than lotic species (Hof et al. 2006, 2012, Grewe et al. 2013, Pinkert et al. 2018). In addition, lentic species markedly dominate in regions that have been covered by glaciers during the last glacial maximum in Europe suggesting a recolonization advantage (Pinkert et al. 2018). However, the drivers of large-scale variation in the richness of freshwater species and the extent to which lentic and lotic species respond to environmental constraints remain largely unknown (reviewed in Pinkert et al. 2022b, but see Acquah-Lamptey et al. 2020).

Here we use comprehensive global distributional and habitat data for odonate species (damselflies and dragonflies) to decompose the role of natural selection on physiological traits and the ecological implications of adaptations to habitat stability underpinning the diversity of freshwater insects. According to the HSDH, the greater dispersal ability should provide an advantage for lentic species in climatically less stable habitats. Thus, (I) the proportion of lentic species should be positively related to short-term (seasonality across decades) and long-term (paleoclimatic fluctuations across millions of years) climatic instability as well as proxies of dispersal capability (body and range size) in odonates. Using the explicit mechanistic framework of the MTE as a baseline, we evaluate the relative contribution of temperature versus adaptations to habitat stability in explaining lentic and lotic species richness. The MTE predicts a negative relationship of logarithmic species richness and inverse temperature (1/kT) (Brown et al. 2004) at a slope between –0.60 and – 0.70. However, adaptations to habitat stability as conceptualized in the HSDH would violate the MTE’s assumption of independent variation in body size and abundance with temperature. This integrated framework allows us to directly test whether departures from the theoretically expected slope or intercept under the MTE can be explained by dispersal– stability dynamics as predicted by the HSDH. Integrating the MTE and the HSDH, we expect (II) that temperature is less (decreases with a shallower slope) and climatic seasonality is more important for lentic species richness than for lotic species richness. A higher intercept for assemblages rich in lentic species would suggest that dispersal–stability dynamics (HSDH) elevate richness above the metabolic baseline predicted by MTE. Conversely, a lower intercept could suggest that additional constraints (e.g., dispersal limitation) depress richness despite metabolic expectations. Consequently, the MTE should systematically underpredicting the overall species richness of assemblages with a high proportion of lentic species.

## Method details

### Distribution data

We used the most complete set of distribution data currently available for odonates (data taken from Mähn et al. 2023). This dataset consists of two types of distributional information: expert range maps and ranges obtained through the intersection of occurrence records with the terrestrial ecoregions of the world. In short, expert range maps provided by IUCN.org (IUCN 2021) and Boudot and Kalkman (2015) were intersected with a standard equal-area grid of cells of approximately 100 km × 100 km size (military grid reference system) and then taxonomically harmonized. As some of these range maps were incomplete or entirely missing for some species, expert range maps were extended with ecoregional ranges (Pinkert et al. 2023). These ecoregional ranges were generated by intersecting spatially cleaned and taxonomically harmonized occurrence records from (Sandall et al. 2022, originally downloaded from GBIF.org, 08 July 2021, DOI: https://doi.org/10.15468/dl.tc7q68) outside expert maps with a global terrestrial ecoregions layer (downloaded from OneEarth.org; (Dinerstein et al. 2017)). If a species had at least one occurrence record within an ecoregion, that entire ecoregion was included as part of its potential range, following the assumption that a verified species presence indicates broader ecological suitability within that unit. Although this approach may overestimate actual distribution in some cases, it generally provides a robust and reproducible method for filling gaps in missing expert range maps largely overlapping with both species distribution models and expert range maps (Pinkert et al. 2023). Ecoregional ranges offer a broader definition of species ranges based on ecological characteristics and expert knowledge, but their average sensitivity and precision outperforms conventional methods particularly for species with less than 100 occurrence records (Pinkert et al. 2023). Finally, the expert and ecoregional species ranges were combined and duplicate cell-species combinations, as well as cells that did not have mean annual temperature data (i.e those with >50% water). The final distribution dataset included 5,233 of the 6,402 (83%) currently described odonate species (accessed July 26.2023, Paulson, et al. 2023).

### Habitat Information

We extracted 5,155 individual records of habitat affiliation data for 3,018 species from descriptions provided in 21 literature sources (Table S1). Distribution data were available for 2,932 of these species (46% of all odonates). 1,062 species were categorized as lentic, 1,458 as lotic and 412 that were recorded in both habitat types (e.g. slow floating rivers or oxbows). For assemblage-level analysis of differences in species richness patterns, we counted the number of co-occurring lentic species and habitat generalists as lentic as well as the number of lotic species. The proportion of lentic species (hereafter simply called ‘lenticity’) was calculated as lentic species richness (*n =* 1,474) divided by the total count of species with habitat information per assemblage. We considered generalists as lentic species, because adaptations to lentic habitats (e.g. dedication and frost tolerance) are a derived feature (Corbet 2004, Letsch et al. 2016). The median coverage of habitat information, this is the count of species with habitat and distribution data divided by the count of species with distribution data, was 92% (min: 22%, max: 100%; see Fig. S1). Note that due to the rather high coverage and to maximize completeness of the data for analyses of the overall species richness, we retained information for all assemblages.

### Environmental data

To assess the relative importance of annual versus seasonal environmental drivers for species richness, we used the three variables mean annual temperature, precipitation, and mean annual enhanced vegetation index (EVI) as well as their corresponding seasonality (standard deviation). Climate data were downloaded from chelsa-climate.org (Karger et al. 2017, Amatulli et al. 2018) and EVI data was downloaded from EarthEnv.org (Amatulli et al. 2018). The EVI layer was cropped to the extent of the climate variables with functions of the R-package ‘raster’ (Hijmans et al. 2016). To reduce multicollinearity in multivariate analyses, we summarized annual and seasonal environmental variables into one principal component each using functions of the R-package ‘stats’. The PCA was based on the correlation matrix, as the variables were measured in different units. In addition, we considered mean elevation because of its common effect on species richness (Rahbek et al. 2019) and the standard deviation of mean annual temperature across 10,000 ya intervals of the Pleistocene (Gamisch 2019), as a proxy for the paleoclimatic instability (also used in Pinkert et al. 2025b).

### Phylogenetic autocorrelation

Closely related species are expected to show similarities in traits and trait-environment responses to environmental changes due to their shared origin. However ordinary least square regression models assume an independence of the data points used in the analysis. We therefore accounted for phylogenetic relationships in species-level analysis of the average body size and range size with lenticity as well as between body size and range size, using phylogenetic generalized least squares regressions. The tree used in these analyses (Mähn et al. 2023) was constructed based on taxonomic data and phylogenetic inferences of internal nodes based on Bybee et al. (Bybee et al. 2021) and Letsch et al. (Letsch et al. 2016). Intra-genus relationships were randomly resolved with the function ‘multi2di’ of the R-package ‘phytools’ (Revell 2017). Branch length was calculated using Grafen’s method (Grafen 1989). Corresponding functions are included in the R-package ‘ape’ (Paradis et al. 2004). The tree was pruned to contain species with information on habitat preference. In addition, we used D statistics (Fritz and Purvis 2010) based on this tree with functions of the R package ‘caper’. D is a measure specifically for binary traits, to calculate the phylogenetic signal in habitat preferences across odonate species.

### Spatial autocorrelation

Spatial autocorrelation in the residuals of regression models is a common bias in macroecological analyses (Dormann et al. 2007). This phenomenon of non-independence among neighbouring grid cells can lead to an overestimation of degrees of freedom and consequently result in erroneous parameter estimates and model inference. To address the issue of spatial autocorrelation and to ensure robust analyses, we assessed the point of spatial independence based on Moran’s *I* using correlograms of the residuals of models and calculating a spatial covariance matrix based on Euclidean distances of the coordinates of assemblages with the point of spatial independence as maximum distance threshold (Fig. S4). These model-specific covariance matrices were integrated into spatial autoregressive models using the R-package ‘spdep’ to obtain model estimates corrected for spatial autocorrelation (Bivand et al. 2017). Subsequent Moran’s I tests of the residuals of these final SAR error models were non-significant (Table 1, Moran’s I ≈ 0, all p = 1), indicating that spatial autocorrelation was effectively accounted for.

**TABLE 1.** Regression models of species richness and the proportion of lentic species in odonate assemblages. Assemblage-level multiple least squares regressions of overall species richness, lentic and lotic species richness, respectively, as well as the proportion of lentic species (lenticity) with environmental variables, including a spatial dependency weight (spatial autoregressive model, SAR). Species richness refers to the total count of odonate species, based on an assessment of 5,233 species with distribution data. Species with habitat preference data were divided into lentic species (*n =* 1,474 species and 18,082 assemblages) and lotic species (*n =* 1,458 species and 16,653 assemblages).

| Dependent variable | Predictor | Estimate | SE | *t*-value | *P* | *R*<sup>2</sup> |
| --- | --- | --- | --- | --- | --- | --- |
| Species richness | Annual PC | 72 | ±0.81 | 88.14 | <0.001 | 0.52 |
|  | Seasonal PC | −2.7 | ±0.70 | −3.75 | <0.001 |  |
| | Elevation [$\log_{10}$] | 22 | ±0.71 | 31.05 | <0.001 | |
|  | Pleistocene Temp. SD | 18 | ±0.77 | 23.13 | <0.001 |  |
| Lentic richness | Annual PC | 33 | ±0.45 | 74.92 | <0.001 | 0.47 |
|  | Seasonal PC | −2.2 | ±0.39 | −5.64 | <0.001 |  |
| | Elevation [$\log_{10}$] | 11 | ±0.39 | 28.66 | <0.001 | |
|  | Pleistocene Temp. SD | 13 | ±0.42 | 29.96 | <0.001 |  |
| Lotic richness | Annual PC | 24 | 0.39 | 62.07 | <0.001 | 0.37 |
|  | Seasonal PC | −0.43 | 0.34 | −1.25 | 0.21 |  |
| | Elevation [$\log_{10}$] | 11 | 0.35 | 32.15 | <0.001 | |
|  | Pleistocene Temp. SD | 4.0 | 0.37 | 10.69 | <0.001 |  |
| Lenticity | Annual PC | −0.069 | 0.0012 | −58.30 | <0.001 | 0.37 |
|  | Seasonal PC | 0.013 | 0.0010 | 12.38 | <0.001 |  |
| | Elevation [$\log_{10}$] | −0.046 | 0.0011 | −44.20 | <0.001 | |
|  | Pleistocene Temp. SD | −0.0036 | 0.0011 | −3.167 | 0.0015 |  |

PC: Principal Component; SD: Standard deviation

### Statistical analyses

Statistical analyses were performed in R (R Core Team, 2023) and the frequency distributions of all variables were visually inspected prior to modelling. Body size and range size were log_10_-transformed to normalize the data. For comparability across models, all predictor variables were z-scaled.

To investigate the drivers of lenticity (i.e. the proportion of lentic species), we fitted both ordinary least squares as well as spatial autoregressive models that account for spatial autocorrelation with the first PC of annual and the first PC of seasonal climatic variables as well as paleoclimatic instability and elevation as predictors (Table S2). Because the proportion of lentic species is bound to values from 0 to 1, we performed a beta regression model. This supplementary analysis confirmed that our initial models adequately represent the limits of variation in lenticity (Table S3).

To explore spatial variation in the slope, significance and explained variance of the six putative drivers we used geographically weighted regressions, including the PCs of annual and seasonal climate, elevation, Pleistocene temperature instability, mean body length and range size as predictors. In contrast to traditional regressions that assume constant linear relationships across the study region, geographically weighted regressions assess spatial variation in model parameters. We expected this particularly relevant for two predictors of lenticity. Firstly, the relationship of lenticity with the first PC of annual climatic variables (mostly mean annual temperature) might be non-stationary, as lentic species should be adapted to both extremely high and low temperatures. Hence lenticity should increase with decreasing temperature in higher latitudes and increase with increasing temperature in lower latitudes. Secondly, if the dynamic history of the northern hemisphere favoured the colonization by lentic species (Pinkert et al. 2018) the presumed positive effect of Pleistocene temperature (paleoclimatic) instability could be restricted to regions with very strong paleoclimatic changes in higher latitudes. Corresponding functions for bandwidth estimation (in our cases a Gaussian kernel with a fixed bandwidth) and geographically weighted regressions are provided in the R-package ‘spgwr’ (Bivand et al. 2023).

To assess the relative importance of annual drivers versus seasonal climate and paleoclimatic instability for the overall species richness of odonates (all 5,233 species with distribution data), lentic species richness (1,474 species) and lotic species richness (1,458 species), we fitted ordinary least squares regressions as well as spatial autoregressive models (Table 1). These models also included elevation as a potential additional predictor of species richness.

To test the predictions of the metabolic theory of ecology (MTE), which suggests that species richness increases exponentially with temperature due to its impact on metabolic rates, we conducted analyses on three different data sets, overall species richness, lentic species richness, and lotic species richness. For each of these sets linear (ordinary least squares) regressions of logarithmic species richness and inverse temperature (1/kT), where *T* is the temperature expressed in Kelvin, and *k* is the Boltzmann’s constant of 8.62 × 10^−5^ eV (Allen et al. 2002, Brown et al. 2004).

Our main analyses considered the observed slope as estimated by an ordinary least squares regression (called ‘MTE-naive model’ hereafter). However, as most species without distribution data are likely missing in regions with higher temperatures (tropical climates), the observed species richness-temperature slope should be shallower than in reality. For analysis of residual variation around the temperature-species richness relationship, we therefore also provide results for a range of three slopes of –0.60, –0.65, and –0.70, suggested in literature (called ‘theoretical MTE models’ hereafter) (Allen et al. 2002, Brown et al. 2004) and analysed their residual variation. These slopes represent the average activation energy of metabolism (∼0.6–0.7 eV for most organisms). Our results confirm that incomplete distribution data for tropical species seems to cause an underestimation of the species richness at high temperatures, flattening the observed log-richness–temperature slope (Fig. 4A). A ddressing this potential sampling bias, we also examined the residual variation around temperature-richness relationships using theoretical MTE slopes (–0.60 to –0.70). These analyses yielded results highly consistent with those based on the (observed) general MTE mode slope, suggesting our main conclusions are robust (Fig. S5). Interestingly, residual variation explained by other environmental variables, such as seasonality, appeared even stronger under these theoretical slopes, further supporting their ecological relevance.

These findings also imply that the variance explained by temperature alone may be underestimated due to the data limitations. In addition, previous studies have emphasized the need to account for differences in the abundance and body size of assemblages as the MTE assumes that these factors are constant. However, these factors are typically not included in larger-scale analysis, neither as control nor as factor of interest. One reason for this limitation is the lack of such data at scale. Here we used body size data taken from a previous study and range size estimates derived from species range maps (i.e. the number of equal-area cells). Range size estimates served as a proxy for assemblage-level species abundance based on the ubiquitous distribution-abundance relationship (Borregaard and Rahbek 2010, Friess et al. 2017, Pinkert et al. 2020a).

## Results

### Environmental predictors of species richness

Overall species richness as well as lentic and lotic odonate species richness broadly increased with decreasing latitude (species richness: estimate ± SE = –1.44 ± 0.021, p < 0.0001, R^2^ = 0.20; lentic species richness: estimate ± SE = –0.64 ± 0.011, p < 0.0001, R^2^ = 0.15; lotic species richness: estimate ± SE = –0.53 ± 0.010, p < 0.0001, R^2^ = 0.13; Fig. 1A). In models that account for spatial autocorrelation, the principal component of annual climate variables consistently had the strongest positive effect on species richness with standardized effect sizes twice as strong as those of other predictors (Table 1). Overall species richness as well as lentic and lotic species richness generally increased with increasing elevation. However, lentic species richness increased with increasing climate seasonality and Pleistocene temperature instability, while lotic species richness was not affected by these two predictors. Together annual climate, climate seasonality, Pleistocene temperature instability and elevation explained 52% and 47% of the variation in overall species richness and lentic species richness, but only 37% of the variation in lotic species richness (Table 1).

**FIGURE 1.**
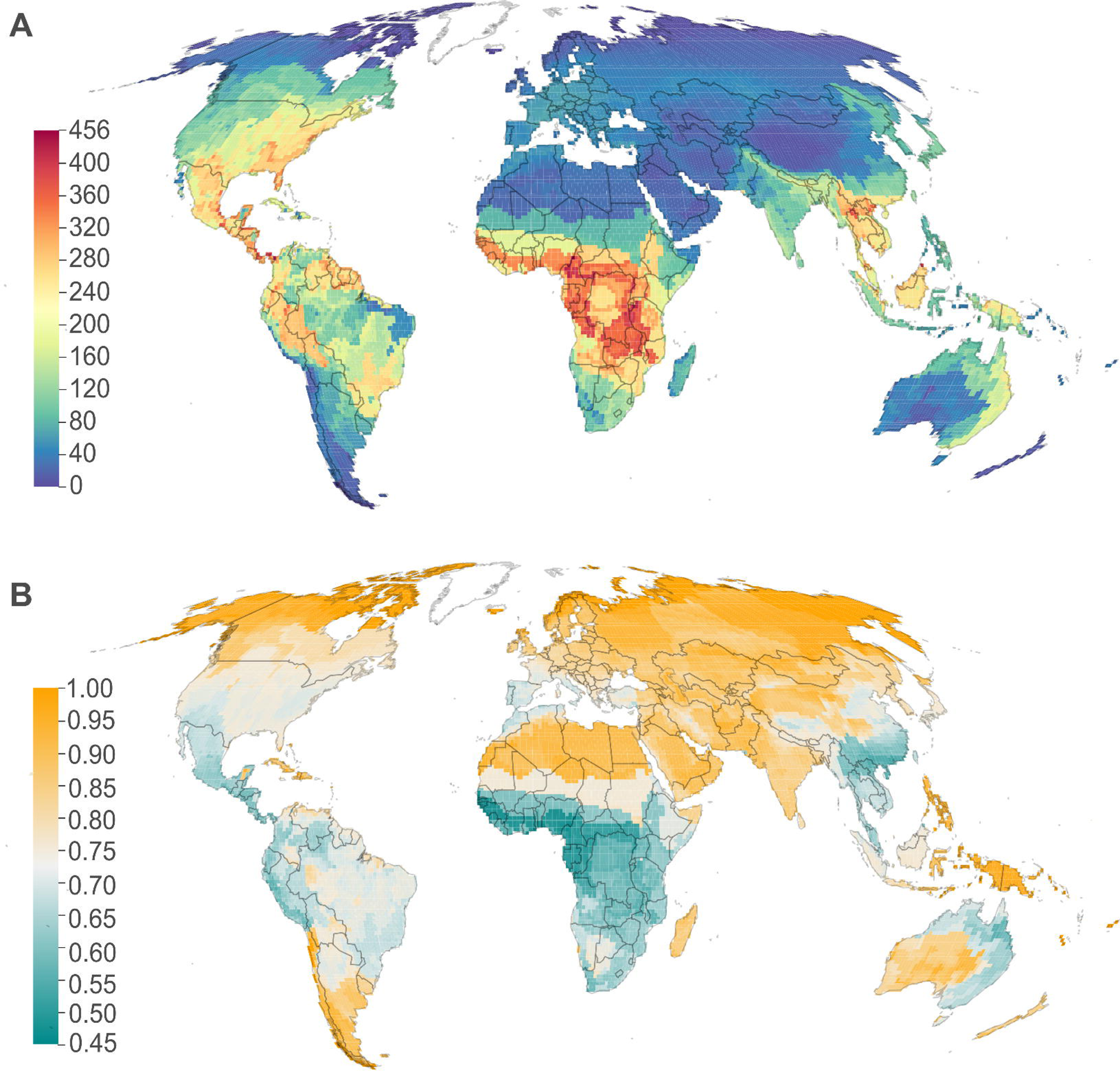
Spatial variation of the overall species richness and the proportion of lentic species. Overall count of odonate species (A, 83% [5,233] of all odonate species) and the proportion of lentic (standing water) species relative to the count of all species with habitat information (B, 46% [*n =* 2,932] of all odonate species) per assemblage (*n* = 18,082 assemblages). Maps are shown in a Mollweide projection. Colour scale intervals follow an equal-class breath classification.

Kernel density estimates across the annual and seasonal climate space confirmed that lentic and lotic species occupy distinctly different environmental conditions. Lentic species mainly clustered at the lower end of annual temperature, productivity, and precipitation gradient (PC1, explained variance 47.2%) and the upper end of seasonality in temperature, precipitation, and productivity (PC2, explained variance 32.3%), whereas lotic species occupy a narrower range at the upper annual and lower seasonal end of the climate space (Fig. S3).

### Drivers of the proportion of lentic species

To better understand differences in the patterns of species richness between lentic and lotic species, we calculated the proportion (%) of lentic species relative to the total count of species with habitat information per assemblage (‘lenticity’). The number of species associated with lentic was similar to that associated with lotic habitats in our data (1,474 versus 1,458 species) and none of the considered assemblages was exclusively composed of lotic species. In turn, a relatively large number of assemblages (1,540 cells, 8.5%) was exclusively composed of lentic species, with lenticity globally being 77% on average (min: 42%, max: 100%). Linear models as well as phylogenetic generalised least squares (pgls) models, where we accounted for the phylogenetic relationship of species, showed that this skew in the composition of odonate assemblages is a direct implication of the on average greater range size of lentic species compared to lotic species (linear model: estimate ± SE = 0.49 ± 0.04, p < 0.001, R^2^ = 0.06; pgls: estimate ± SE = 0.40 ± 0.07, p < 0.001, R^2^ = 0.02).

Lenticity generally increased with increasing latitude, and was lowest in South Africa, Southeast Asia, central Africa, Mesoamerica and the Andes (Fig. 1B). Compared to species richness patterns of the two groups, effects of annual estimates on lenticity were weaker and explained less variance (Table S1). All environmental predictors together explained 56% of the variation in lenticity in models accounting for spatial autocorrelation (Table S3). The average body size (Mähn et al. 2023) of assemblages decreased and the average range size increased with increasing lenticity. Also, lenticity showed a strong latitudinal gradient with an increasing proportion of lentic species towards the poles with absolute latitude explaining 22% of this variation (linear model: estimate ± SE = 1.69 ± 0.023, p < 0.001, R² = 0.22).

Lentic species were on average smaller (linear model of log_10_(body length): estimate ± SE = –0.378 ± 0.043, p < 0.0001, R^2^ = 0.04) and had larger ranges than lotic species (linear model of log_10_(range size): estimate ± SE = 0.772 ± 0.060, p < 0.0001, R^2^ = 0.06). Also, a smaller body size was associated with larger range sizes in odonates (both log_10_-transformed; linear model: –0.079 ± 0.019, p < 0.0001, R^2^ = 0.01). However, the variance explained by these models was small when controlling for the phylogenetic relationship of species (pgls for body size∼habitat, range size∼habitat and body size∼range size: estimate ± SE = 0.099 ± 0.053,0.039 ± 0.066, – 0.045 ± 0.017; p = 0.19, < 0.001, 0.010; R^2^ < 0.01, 0.02, < 0.01). While their significance underscores robust global relationships, these very minor effects reflect the complexity and variability of environmental drivers across various geographical regions inherent to global-scale analyses at the species-level (Fig. Appendix S5). By specifically addressing spatial trait variation through assemblage-level analyses, and hence at the level of biological organisation most relevant for geographical variation in species richness, the same effects were orders of magnitude stronger (Table S3; see also Laumeier et al. (2023). Thus, assemblage-level lenticity was also positively but more strongly related to body size and range size (linear models: estimate ± SE = 1.86 ± 0.38 and 1.22 ± 0.02, both p < 0.001, R^2^ = 0.13% and 12%, respectively) than for species-level differences.

Species-level phylogenetic analysis confirmed a strong signal of evolutionary conservatism (D statistic = 0.32) in the habitat preference of odonates. Lentic species had their range centre generally at higher (absolute) latitudes than lotic species (linear model: estimate ± SE = 5.01 ± 0.57, p < 0. 001, R^2^ = 0. 03). Of all 39 considered odonate families, 19 were exclusively lotic and 2 exclusively lentic according to our data. However, only 13 families had sufficient data (<5 lentic and 5 lotic, but at least 10 species in total) to assess differences regarding the latitudinal range centre of species within families with respect to habitat preference. In six of these families, comprising in total 1966 species (67% of all species in our data), lentic species had range centres in higher latitudes (absolute) than lotic species (Fig. 3). The only family that showed an opposite trend was Calopterygidae, comprising 13 species.

Given the expected non-stationarity of the effects of some predictors of lenticity, we also fitted geographically weighted regressions. These analyses showed minor variation in the effects of annual climate variables and elevation, with overall negative effects on lenticity (Fig. 2). Seasonal climate variables and range size generally positively affected lenticity. However, the effects of Pleistocene temperature stability and average body size were highly localized. Body size negatively affected lenticity in subtropical parts across South America, Africa and Eurasia and strongly positively affected lenticity in western North America, central Africa, South Africa, and Australia. The effects of seasonal temperature, productivity and precipitation (Seasonal PC) were positive in temperate but negative in tropical regions. Elevation generally had a positive effect largely limited to higher elevations across the globe. Paleoclimatic instability had a highly local positive effect on lenticity in higher latitudes of the northern hemisphere. The main trends in geographically weighted regressions of lenticity are underscored by results from single and multiple regressions including the same predictors (Table S1). However, while Pleistocene temperature instability had a strong positive effect in single regressions and alone explains 11% of the variation in lenticity, it had no effect in spatial autoregressive models including all six potential predictors (Table 1).

**FIGURE 2.**
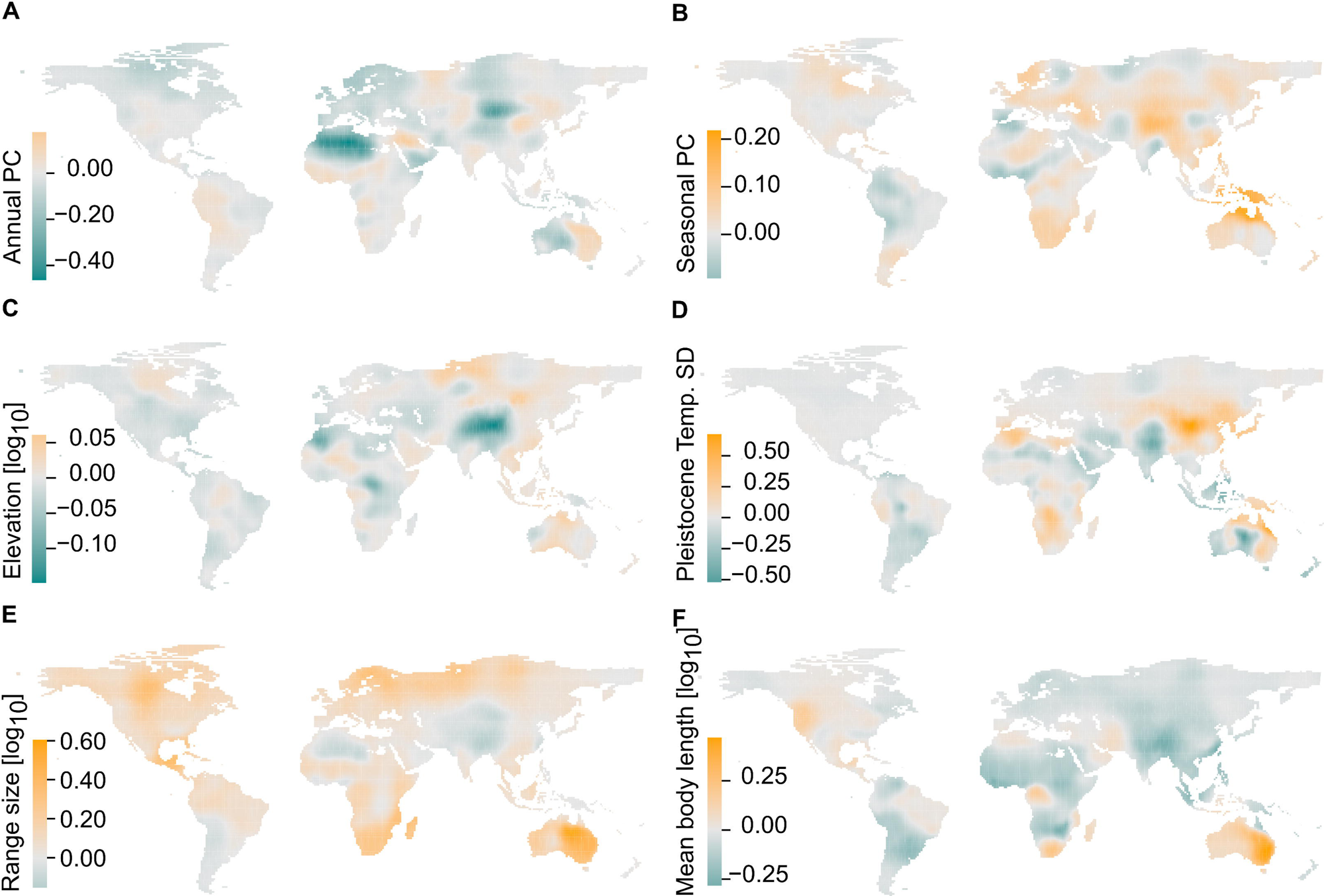
Spatial variation in standardized coefficients of predictors of lenticity from geographically weighted multiple regression. The maps represent local relationships of the lenticity of 18,082 odonate assemblages with (A) the principal component of annual climatic variables, (B) the principal component of seasonal climatic variables, (C) log_10_-transformed elevation, (D) the Pleistocene temperature stability (standard deviation of annual mean temperature across the Pleistocene), (E) log_10_-transformed average range size and (F) log_10_-transformed average body length. Maps are shown in a Mollweide projection. Colour scale intervals follow an equal-class breath classification.

### Testing the MTE predictions and assessing drivers of residual variation

We tested the MTE predictions by comparing the observed slope from the ordinary least squares regression (‘general MTE model’) with theoretically derived slopes from the literature (‘specific MTE model’; Allen et al. 2002, Brown et al. 2004). These slopes are theoretically inferred expectations under the assumption of uniform metabolic scaling and energy constraints across taxa. The observed slope for overall odonate species richness was – 0.34 (Fig. 4A), much shallower than the theoretical expectation of –0.60 to –0.70 for ectothermic species (Allen et al. 2002). Temperature alone explained 37% of the variation in overall species richness in both regressions based on this observed slope and a specific slope of –0.65 (Fig 4A). When distinguishing species of the two habitat types, lotic assemblages showed a steeper (–0.47, R^2^ = 0.24) slope than lentic assemblages (–0.28, R^2^ = 0.32; Fig. 4B). Notably, assemblages with both high and low proportions of lentic species were found at the warm end of the temperature gradient (below 0° C or approximately 42.5 inverse temperature), with lentic assemblages comprising substantially fewer species than lotic assemblages here. By contrast, assemblages found at the cold end of the temperature range were almost exclusively composed of lentic species (Fig. 4D). Lenticity explained 33% and 38% (21% and 24% absolute variance; Spearman’s rho = –0.59; Fig. 4D) of the residual species richness variation from the general MTE model in ordinary least squares and spline-based smoothed regressions, respectively. This residual variation was generally negatively associated with lenticity, with more negative deviation in cold and more positive deviations at the warm end of the gradient.

Regions with less species than predicted by the general MTE model (Fig. 4C) highlighted the Sahara region, the Middle East, Madagascar, the Atacama, and central Australia, as well as polar regions in the northern hemisphere, Tierra del Fuego and the Tibetan highlands. Regions with more species than predicted highlighted the East Coast of North America, Central African mountain ranges, the Suntar-Khayata range, southern slopes of the Himalaya and the northern Andes.

The results of our general MTE model (i.e. observed slope of –0.34) that tested for predictors of species richness, beyond inverse temperature, showed that species richness was strongly underpredicted in regions where annual productivity, annual precipitation (PC of annual climatic predictors) and assemblage-level average body size was higher (Fig. 4C). By contrast the effect of seasonal climatic predictors was comparatively low. However, in models that are fitted with theoretical MTE slopes (–0.60, –0.65 or –0.70), the effect of additional annual climatic drivers was reduced to a minimum and the effects of average body size were weaker, while those of seasonal climatic predictors were substantially greater. This indicates that models which do not account for the temperature-specific assumptions of the MTE systematically underestimate the contribution of seasonal climate and overestimate that of productivity and other annual climatic predictors.

## Discussion

Our results provide insights into the environmental drivers of dragon- and damselfly species richness and exemplify the crucial role of adaptations to spatial and temporal habitat stability for promoting the diversity of freshwater systems. With this first global scale analysis of the environmental drivers of species richness for an entire order of freshwater insects, we rigorously underscore the strong, general increase in species richness with increasing temperature, productivity, but also the influence of climate seasonality associated with species’ habitat preference that had broad impacts on the biogeographical distribution and history of odonate lineages.

Consistent with previous studies on a wide range of organisms (Currie 1991, Hawkins et al. 2003) as well as realm-level diversity gradients in Odonata (Pinkert et al. 2018, 2020b, Abbott et al. 2022), we demonstrate that overall odonate species richness, and the richness of both lentic and lotic odonate species, generally increases with decreasing latitude. Strong impacts of mean annual temperature, productivity and precipitation underpin this latitudinal diversity gradient highlighting the dominant role of constraints in resource availability(Hawkins et al. 2003). With explaining 37% of the variation in overall species richness alone, temperature was the single most important driver of the global species richness of odonates and our results broadly underscore the predictive power of Metabolic Theory of Ecology (MTE). Compared to previous evidence for the MTE across taxa, the strength of the temperature-species richness relationship observed in odonates and the variance in species richness explained by temperature stand out as remarkably strong global support (Hawkins et al. 2007). Moreover, by integrating predictions of the HSDH on adaptations to climate seasonality and adaptations of lentic species that are also linked to the MTE through assumptions of stationary in energy use across space (body size and range size as a proxy for abundance, Cassemiro and Diniz-Filho (2010), our results show that a substantial part of the deviations from the MTE are related to the HSDH trait syndrome. Thus, the proportion of lentic species alone explained 33-38% of the residuals variance from the MTE relationship (Fig. 4). In addition, we found that the intercept of the MTE relationship for lentic species was consistently higher than for lotic species. The higher intercept indicates that lentic assemblages maintain greater baseline richness than predicted by temperature alone, suggesting that dispersal–stability dynamics confer an additive richness advantage. This implies that even at equivalent thermal conditions, lentic assemblages are systematically richer, likely due to their enhanced colonization ability, broader range sizes, and evolutionary history of diversification in unstable habitats.

Beyond improving predictions of species richness, it is crucial to understand idiosyncrasies in temperature-responses among ecological and taxonomic entities that structure communities, because both spatial variation in body size and range size are emergent patterns of biogeographical processes (Pinkert et al. 2020a, Mähn et al. 2023). As conceptualized in the habitat-stability-dispersal hypothesis (Southwood 1977), greater dispersal propensity of lentic compared to lotic species themselves should have evolved as an adaptation to the lower spatial and temporal stability of their habitats. Our global scale test of the HSDH confirms that both climate seasonality and paleoclimatic instability are of far greater importance in driving lentic compared to lotic species richness and largely determine the disproportionate richness of lentic species in higher latitudes. Our results also reconcile smaller-scaled evidence of major ecological differences related to habitat preferences, with overall stronger environment-body size relationships, a smaller body size (Acquah - Lamptey et al. 2020), and larger ranges of lentic compared to lotic species (Hof et al. 2006). Broad differences in the reaction norm are further underscored by regional-scale studies across North America and Europe for odonates, caddisflies, and water beetles, showing that lentic species tend to exhibit greater dispersal ability and generally larger range sizes than lotic species (Hof et al. 2006, 2012; Grewe et al. 2013; Pinkert et al. 2018). Incorporating factors related to these differences of lentic and lotic species into models of species richness explained 45% of the residual species richness variation from the observed (general MTE) and as much as 56% residual variation from models with theoretical MTE slopes (Fig. S5). This showcases the potential of merging physiological and dispersal - related predictions to resolve ambiguity in previous results (Algar et al. 2007, Cassemiro and Diniz-Filho 2010).

A key finding of our study is the phylogenetic conservatism in habitat preference among odonates that seems to have promoted the diversification and distributional success of lentic lineages. Across all analyses, lentic odonate species consistently exhibited higher dispersal ability, broader range sizes and more poleward range centres than lotic species Our results extent earlier regional findings suggesting that lentic species tend to have range centres at higher latitudes and greater dispersal abilities than lotic species (Hof et al. 2006, Marten et al. 2006, Abbott et al. 2022) to the global scale. Lentic species occur in 20 of 39 odonate families, and dominate in all major lineages of odonates found in higher latitudes. Among the 13 families with sufficient numbers of both types of habitats, lentic species’ showed higher absolute latitude range centres than lotic species in six families (i.e. 1966 species - 67% of all species in our data). These families included, for instance Coenagrionidae and Libellulidae, the most species-rich families globally, which together comprise approximately one-third of all odonates (Fig. 3) (Kalkman et al. 2008, Sandall et al. 2022). Particularly, assemblages in cold climates and those regions that were repeatedly glaciated during the Pleistocene are almost exclusively composed of lentic species. Pleistocene temperature instability generally positively affected lenticity and explained a substantial part of its variation (R^2^ = 0.11). Geographically weighted models also highlight a highly regional impact at higher latitudes both in the northern and southern hemisphere (Fig. 2). Together these findings provide global support for a strong historical signature of past colonization events with broad impacts on the phylogenetic and functional composition of odonate assemblages (Pinkert et al. 2018).

**FIGURE 3.**
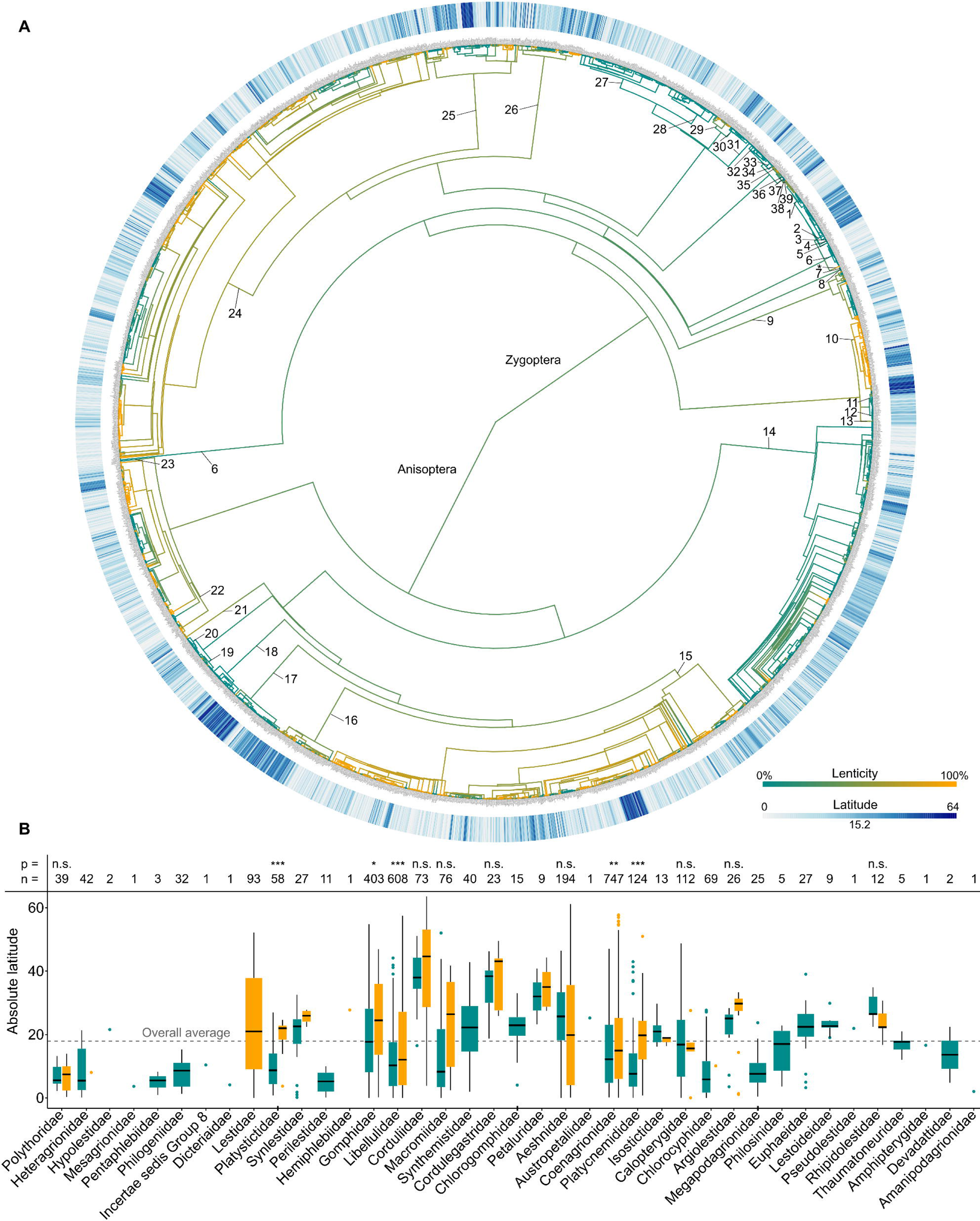
Ancestral trait reconstruction of the habitat affiliation. (A) Brownian motion model of trait evolution (branch and node colours) for the habitat preference of 2,932 (46% of all) odonate species. (B) Average absolute species’ range latitude (see also colour-ring in A) per family for lentic (orange) and lotic (cyan) species. Statistics above the bars are from a multiple linear model including an interaction term of habitat and family as predictor. Asterisks indicate significant differences within family with at least five lentic and five lotic species (p-values: * < 0.05, ** < 0.01, *** < 0.001, n.s. = not significant; *n* = species per family). Families in B are ordered according to their phylogenetic position (numbers in A).

While the Metabolic Theory of Ecology has been tested primarily in terrestrial organisms, its physiological foundations facilitates a general understanding of biodiversity patterns across realms, scales, and taxonomic groups (Allen et al. 2002, Brown et al. 2004, Ruggiero and Hawkins 2006). Our findings shed light on the limitations of applying the MTE to freshwater ecosystems, but also inform about general improvements for predictions of species richness and forecasting responses to climate change. Specifically, we demonstrate that incorporating additional factors, such as habitat stability and dispersal dynamics, is essential to fully leverage the predictive potential of the Metabolic Theory of Ecology (MTE). Thus, alone by accounting for differences in species’ habitat preferences, the explained variance in species richness increased from 37% to 61% in our models (Fig. 4A/D).

**FIGURE 4.**
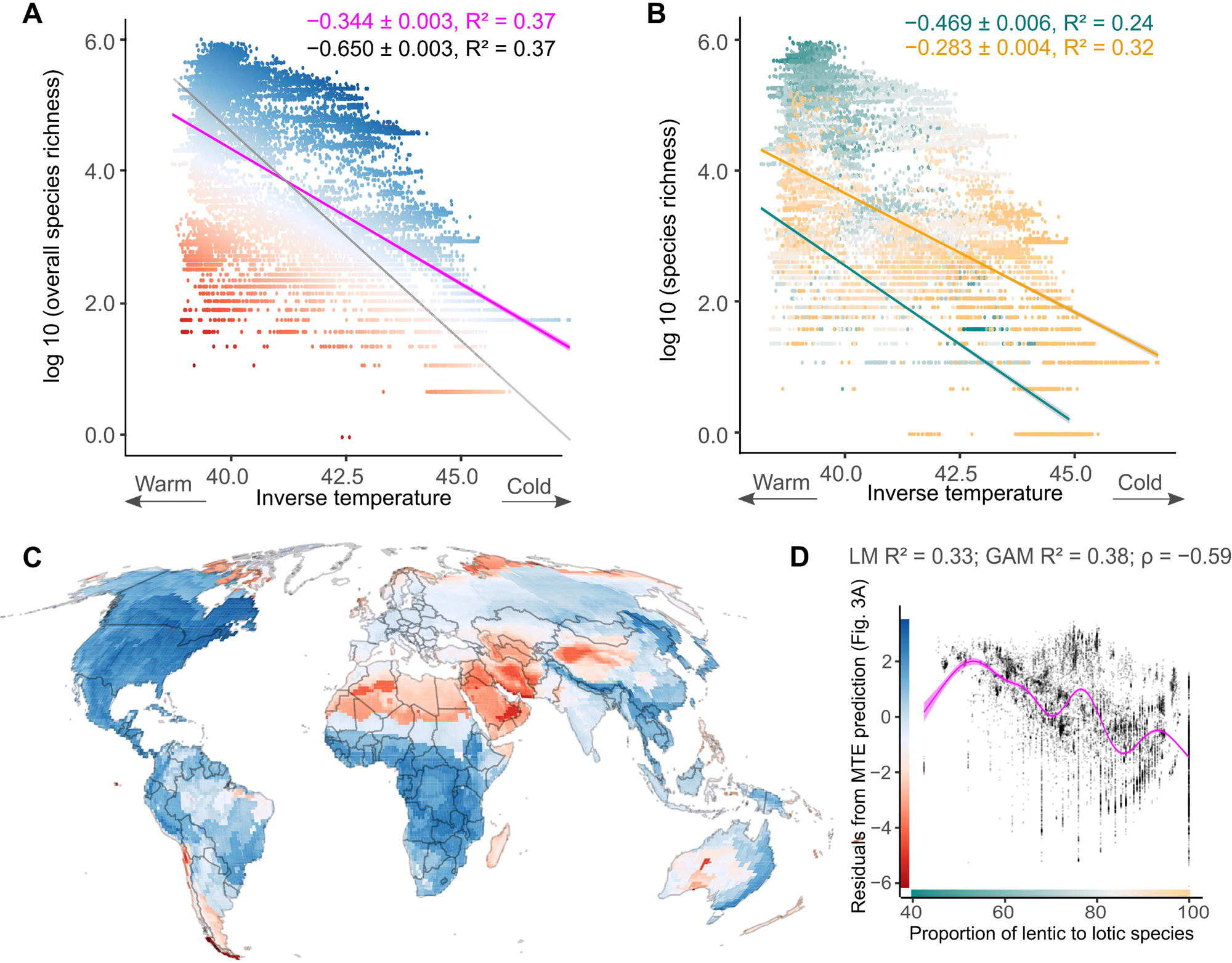
Metabolic scaling for the species richness of odonate assemblages and its residual variation. (A) Relationship of log_10_ transformed species richness and inverse mean annual temperature of 18,082 odonate assemblages. Model statistics and regression lines are provided for both a model fitted with the theoretical slope of –0.65 (grey) as predicted by the Metabolic Theory of Ecology (MTE) and a MTE-naive model (i.e. observed slope, magenta). Regions with more species than predicted are indicated in blue, while regions with fewer species than predicted are marked red. (B) Relationship of log_10_-transformed species richness and inverse mean annual temperature of assemblages of lentic (orange) and assemblages of lotic species (cyan). (C) Spatial variation of the residuals from the MTE-naive regression between overall species richness and inverse temperature (Mollweide projection). (D) Relationship of the residuals from the MTE-naive regression between overall species richness and inverse temperature with the proportion of lentic species (log_10_-transformed). Point colours in A represent the residuals variation (y-axis in D and grid colours in C) and those in B represent the proportion of lentic to lotic species per assemblage (min = 40%, max = 100%, x-axis in D).

Due to missing distribution data in the tropics, biased estimates of species richness at the warm end of the MTE relationship likely resulted in a shallower slope of the observed relationship (Fig. 4A). Notably our trait data is based on an unbiased set of species per literature source allowing comparison of the slopes of lentic and lotic species richness and interpretations of the effect of the proportion of lentic species on the residuals from the overall species richness temperature relationship. However, analysis of the environmental drivers of the MTE model residuals could be sensitive to the spatial bias in data availability. Thus, by additionally analysing the residuals from species richness-temperature models with theoretical MTE slopes, we assessed the robustness of our results regarding the potential effects of this sampling bias. Showing full agreement with the effects of environmental drivers, body size and range size with the MTE-naïve models, our conclusions appear to be robust to a sampling bias. Importantly, this supplementary analysis revealed consistently greater effects of climatic seasonality suggesting an even greater role of habitat stability and improvements beyond the already exceptionally high variance explained by temperature (R^2^ = 37% c.f. 16% across 23 regional studies [Hawkins et al., 2007]), with complete distribution data. Moreover, carefully integrating the assumptions of the MTE, particularly the expected slopes, may inform about the location of poorly sampled regions.

## Conclusions

In conclusion our results emphasize that the potential of the MTE framework for predicting species richness in space and time also extends to freshwater diversity, but cautious implementation of assumptions on spatial invariance of the relationships between body size, and proxies of species’ abundance are crucial to leverage its full potential. These additional factors are also indicative of important idiosyncrasies of species adapted to habitats with broadly different spatio-temporal stability. Thus, despite of the relative stability of lotic habitats, weak dispersal and a greater importance of physiological requirements likely render lotic species more susceptible to the impacts of climate change (Clausnitzer et al. 2009, Dijkstra et al. 2014). Indeed, lentic species have been observed to exhibit stronger northward range expansions (Grewe et al. 2013), lesser abundance declines (Powney et al. 2015, Bowler et al. 2021, Pinkert et al. 2022a), and as well as lower threat levels compared to their lotic counterparts (Rocha-Ortega et al. 2022). Such adaptations to habitat stability are not limited to freshwater organisms but also appear to result in idiosyncratic species richness-temperature relationships of burrowing compared to non-burrowing mammals (Pinkert et al. 2025b). Indeed, despite being mainly applied to freshwater systems in recent years, the habitat-stability-dispersal hypothesis represents a far more general concept with broad implications for understanding the role of spatial and temporal stability of environmental conditions (Southwood 1977). Beyond an improved capability to predict and model community-level diversity, integrating a trait-based ecological perspective into applications of the MTE framework therefore offers an improved understanding of biological responses to environmental changes.

## Acknowledgements

We thank Klaas-Douwe Dijkstra for helpful comments on earlier versions of the manuscript. We acknowledge financial support by the German Research Foundation (grant 409487552) to S.P., R.B and C.H and by the Alexander-von-Humboldt Foundation to S.P.. S.B. also acknowledges support by the National Science Foundation through (grant DEB-2002432). Support for Open Access funding is enabled by the Phillipps-Universität Marburg and organized by the Project DEAL.

## Author Contributions

Conceptualization: S.P., R.B.; Resources: L.A.M.; Data curation: L.A.M.; Formal analysis: L.A.M.; Supervision: S.P.; Funding acquisition: S.P; R.B., C.H., S.B.; Validation: L.A.M., S.P.; Investigation: L.A.M.; Visualization: L.A.M., S.P.; Methodology: L.A.M., S.P., R.B., C.H.; Writing - original draft: L.A.M, S.P.; Project administration: S.P., R.B.; Writing - review & editing: L.A.M., S.P. R.B., C.H., S.B.

