## Appendix for "Metabolic ecology and habitat stability explain the disproportionately high species richness in standing waters"

Ecological Monographs

**Appendix S1**

### Metabolic ecology and habitat stability explain the disproportionately high species richness in standing waters

Laura Anna Mähn, Christian Hof, Seth Bybee, Roland Brandl, Stefan Pinkert

Content:

Table S1-S5

Figure S1-S5

Supplementary references

### TABLE S1 Literature sources of habitat preference data.

| Continent | Title | Reference |
|---|---|---|
| Africa | A Guide to the Dragonflies and Damselflies of South Africa[1] | Tarboton & Tarboton (2019) |
| Africa | The Dragonflies and Damselflies of South Africa[2] | Samways (2008) |
| Asia | Field Guide to the damselflies of New Guinea[3] | Kalkman & Orr (2013) |
| Asia | Field Guide to the dragonflies of New Guinea[4] | Orr & Kalkman (2015) |
| Asia | Dragonflies of Peninsular Malaysia and Singapore[5] | Orr (2005) |
| Asia | The Dragonflies of Sri Lanka[6] | de Fonseka (2000) |
| Asia | Dragonflies and Damselflies of China[7] | Haomiao (2018) |
| Asia | Dragonflies of Russia: Illustrated Photo Guide[8] | Onishko & Kosterin (2021) |
| Asia | Korean Odonata Adult and Larva[9] | Cho (2021) |
| Australia | Dragonflies & Damselflies of New Zealand[10] | Marinov & Ashbee (2020) |
| North America | Damselflies of Texas: A Field Guide[11] | Abbott (2011) |
| Central America | Dragonflies and Damselflies of Costa Rica: A Field Guide[12] | Paulson & Haber (2021) |
| Europe | Field Guide to the Dragonflies of Britain and Europe[13] | Dijkstra & Lewington (2006) |
| North America | Dragonflies & Damselflies of the East[14] | Paulson (2011) |
| North America | Dragonflies & Damselflies of the West[15] | Paulson (2009) |
| South America | Dragonflies of the Colombian Cordillera occidental, A look from the Tatama[16] | Cornelio A Bota-Sierra, Juliana Sandoval-H, Daniela Ayala-Sánchez, & Rodolfo Novelo-Gutiérrez (2019) |
| North America | Damselfly Genera of the New World[17] | Rosser W. Garrison, Natalia von Ellenrieder, Jerry A. Louton (2010) |
| Europe | The dragonflies of europe[18] | R.R. Askew (2004) |
| Asia | Atlas of the dragonflies and damselflies of West and Central Asia[19] | J.-P. Boudot, S. Borisov, G. De Knijf, R. H. A. van Grunsven, A. Schröter & V. J. Kalkman (2021) |
| Australia | The complete field guide to dragonflies of Australia[20] | Theischinger & Hawking (2006) |
| South America | A Guide to the Dragonflies & Damselflies of the Serra dos Órgãos[21] | Tom Kompier (2015) |

**TABLE S2 Ordinary least squares regression models of species richness and the proportion of lentic species in odonate assemblages.** Assemblage-level ordinary least squares regression models between species richness, lentic respectively lotic species richness, as well as the proportion of lentic species and environmental predictors. Results are provided for both single and multiple regressions of these relationships. Species richness refers to the total count of species with distribution data (83% [5,233] of all odonate species), whereas lentic and lotic species richness as well as the proportion of lentic species (lenticity) was calculated based on a subset of (46% [2,932] of all odonate species) species with habitat information (1,474 lentic species and all 18,082 assemblages; 1,458 lotic species and only 16,653 assemblages).

| Dependent variable | Predictor | Estimate | SE | *t*-value | *P* | $R^2$ |
|---|---|---|---|---|---|---|
| Species richness | Annual PC | 68.49 | ±0.52 | 132.2 | <0.001 | 0.49 |
| | Seasonal PC | 2.10 | ±0.73 | 2.88 | 0.004 | 0.00 |
| | Elevation [log] | 6.56 | ±0.91 | 7.20 | <0.001 | 0.00 |
| | Pleisto. temp. SD | –18.45 | ±0.69 | –26.82 | <0.001 | 0.04 |
| Lentic richness | Annual PC | 31.93 | ±0.29 | 108.3 | <0.001 | 0.39 |
| | Seasonal PC | 0.93 | ±0.38 | 2.46 | 0.014 | 0.00 |
| | Elevation [log] | 4.64 | ±0.47 | 9.78 | <0.001 | 0.00 |
| | Pleisto. temp. SD | –5.63 | ±0.36 | –15.5 | <0.001 | 0.013 |
| Lotic richness | Annual PC | 23.91 | ±0.28 | 82.98 | <0.001 | 0.29 |
| | Seasonal PC | –0.93 | ±0.38 | –2.79 | <0.001 | 0.00 |
| | Elevation [log] | 3.93 | ±0.41 | 9.69 | <0.001 | 0.01 |
| | Pleisto. temp. SD | –8.09 | ±0.31 | –26.27 | <0.001 | 0.03 |
| Lenticity | Annual PC | –8.76 | ±0.073 | –119.7 | <0.001 | 0.44 |
| | Seasonal PC | 2.21 | ±0.100 | 22.78 | <0.001 | 0.03 |
| | Elevation [log] | –2.57 | ±0.121 | –21.13 | <0.001 | 0.02 |
| | Pleisto. temp. SD | 4.26 | ±0.089 | 47.83 | <0.001 | 0.11 |
| Species richness | Annual PC | 80.02 | ±0.58 | 138.30 | <0.001 | 0.54 |
| | Seasonal PC | –3.66 | ±0.58 | –6.30 | <0.001 | |
| | Elevation [log] | 22.72 | ±0.64 | 32.23 | <0.001 | |
| | Pleisto. temp. SD | 19.48 | ±0.63 | 31.14 | <0.001 | |

| | | | | | | |
|---|---|---|---|---|---|---|
| | Annual PC | 40.03 | ±0.33 | 123.61 | <0.001 | |
| | Seasonal PC | –3.98 | ±0.33 | –12.26 | <0.001 | |
| Lentic richness | Elevation [log] | 12.92 | ±0.36 | 35.79 | <0.001 | 0.47 |
| | Pleisto. temp. SD | 14.46 | ±0.35 | 41.29 | <0.001 | |
| | Annual PC | 29.43 | ±0.34 | 87.62 | <0.001 | |
| | Seasonal PC | –4.71 | ±0.31 | –15.15 | <0.001 | |
| Lotic richness | Elevation [log] | 11.95 | ±0.35 | 34.44 | <0.001 | 0.35 |
| | Pleisto. temp. SD | 5.66 | ±0.33 | 16.92 | <0.001 | |
| | Annual PC | –9.78 | ±0.079 | –123.00 | <0.001 | |
| | Seasonal PC | 2.18 | ±0.080 | 27.38 | <0.001 | |
| Lenticity | Elevation [log] | –3.96 | ±0.089 | –44.69 | <0.001 | 0.52 |
| | Pleisto. temp. SD | –1.25 | ±0.086 | –14.58 | <0.001 | |

**TABLE S3 Ordinary least squares regression models of the proportion of lentic species in odonate assemblages.** Assemblage-level ordinary least squares regressions and spatial autoregressive models (SAR) of the relationship of the proportion of lentic species with environmental predictors, range size and body size (n = 18,082 assemblages).

| Dependent variable | Predictor | Estimate | SE | *t*/*z*-value | *P* | $R^2$ |
|---|---|---|---|---|---|---|
| Lenticity (LM) | Annual PC | –8.04 | ±0.10 | –87.36 | <0.001 | 0.48 |
| | Seasonal PC | 3.87 | ±0.10 | 40.30 | <0.001 | |
| | Elevation [log] | –3.72 | ±0.10 | –40.67 | <0.001 | |
| | Pleistocene. Temp. SD | –1.02 | ±0.10 | –11.42 | <0.001 | |
| | Mean body length (log) | –3.32 | ±0.10 | –32.66 | <0.001 | |
| | Range Size (log) | 1.87 | ±0.10 | 22.19 | <0.001 | |
| Lenticity (SAR) | Annual PC | –6.18 | ±0.10 | –59.57 | <0.001 | 0.56 |
| | Seasonal PC | 2.63 | ±0.10 | 27.20 | <0.001 | |
| | Elevation [log] | –3.16 | ±0.09 | –36.96 | <0.001 | |
| | Pleistocene Temp. SD | –0.38 | ±0.10 | –3.96 | <0.001 | |
| | Mean body length (log) | –4.29 | ±0.10 | –44.26 | <0.001 | |
| | Range Size (log) | 4.51 | ±0.11 | 39.44 | <0.001 | |

**TABLE S4 Multiple least squares beta regression models of the proportion of lentic species.** Assemblage-level multiple least squares beta regression model of the proportion of lentic species and environmental predictors for 18,082 odonate assemblages.

| Dependent variable | Predictor | Estimate | SE | z-value | *P* | $R^2$ |
|---|---|---|---|---|---|---|
| Lenticity | Annual PC | –0.77 | ±0.91 | –84.32 | <0.001 | 0.31 |
| | Seasonal PC | 0.43 | ±1.80 | 48.39 | <0.001 | |
| | Elevation [log] | –0.47 | ±1.81 | –51.99 | <0.001 | |
| | Pleistocene Temp. SD | –0.11 | ±0.91 | –11.74 | <0.001 | |
| | Mean body length (log) | –0.14 | ±0.83 | –16.11 | <0.001 | |
| | Range Size (log) | 0.16 | ±0.80 | 19.75 | <0.001 | |

**TABLE S5 Principal component analysis loadings for annual and seasonal variables.** Loadings from principal component analysis used for the annual and for the seasonal variable, respectively. Each loading represents the contribution of the original variables to the corresponding principal component.

| Environmental variable | PC Annual | PC Seasonal |
|---|---|---|
| A. Temperature | 0.75 | –0.57 |
| A. Precipitation | 0.84 | 0.36 |
| A. Productivity | 0.88 | 0.31 |
| S. Temperature | –0.09 | 0.30 |
| S. Precipitation | –0.09 | –0.80 |
| S. Productivity | 0.01 | 0.65 |

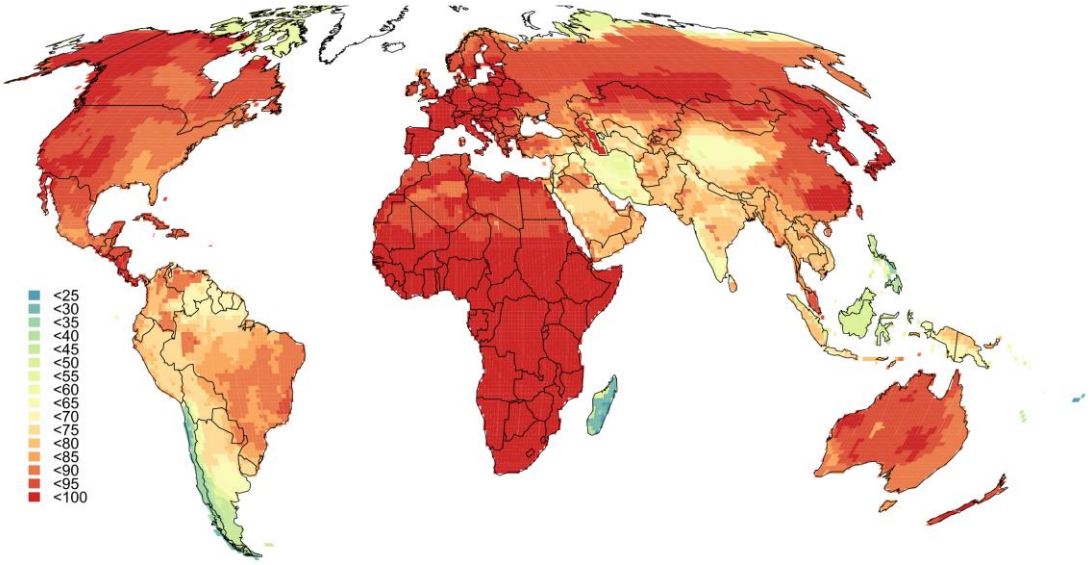


**FIGURE S1 Coverage of habitat information.** Spatial variation in the coverage of species with habitat information per assemblage (18,082 assemblages). Species coverage as the count of species with habitat information (2,932) divided by the count of species for which distributional data was available (5,233; Fig. 1). Colour class intervals are scaled to equally wide value ranges (5% intervals) from 0% (blue) to 100% (red).

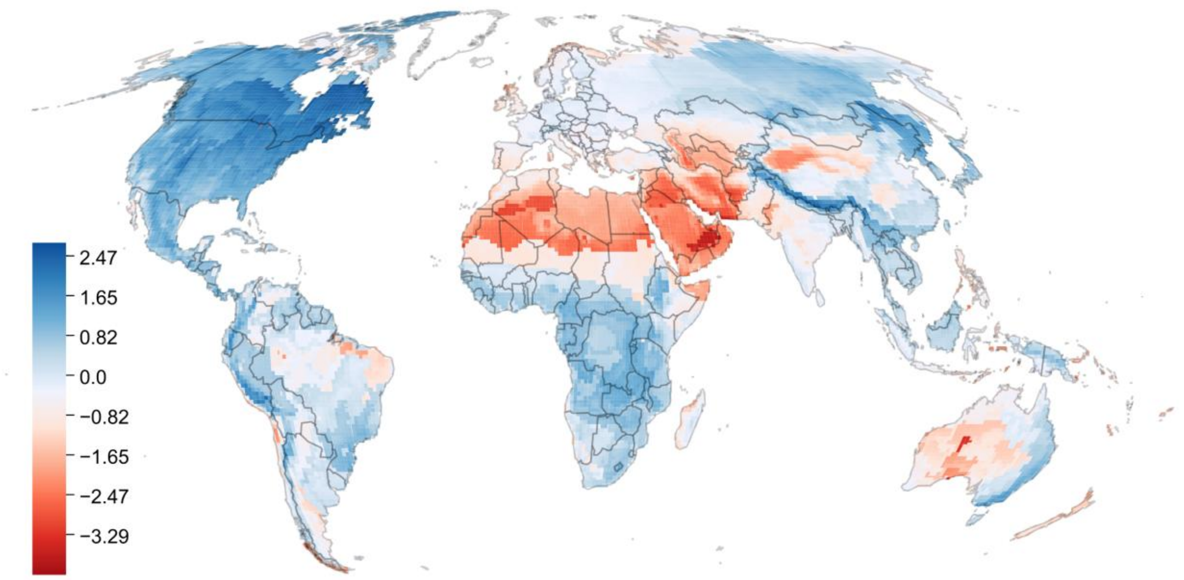


**FIGURE S2 Spatial variation of residuals from the idealized MTE-based prediction.** Residuals are calculated based on the relationship of the overall odonate species richness of 18,082 odonate assemblages and inverse temperature fitted with the idealized slope of –0.65 as predicted by the Metabolic Theory of Ecology. The map is shown in a Mollweide projection.

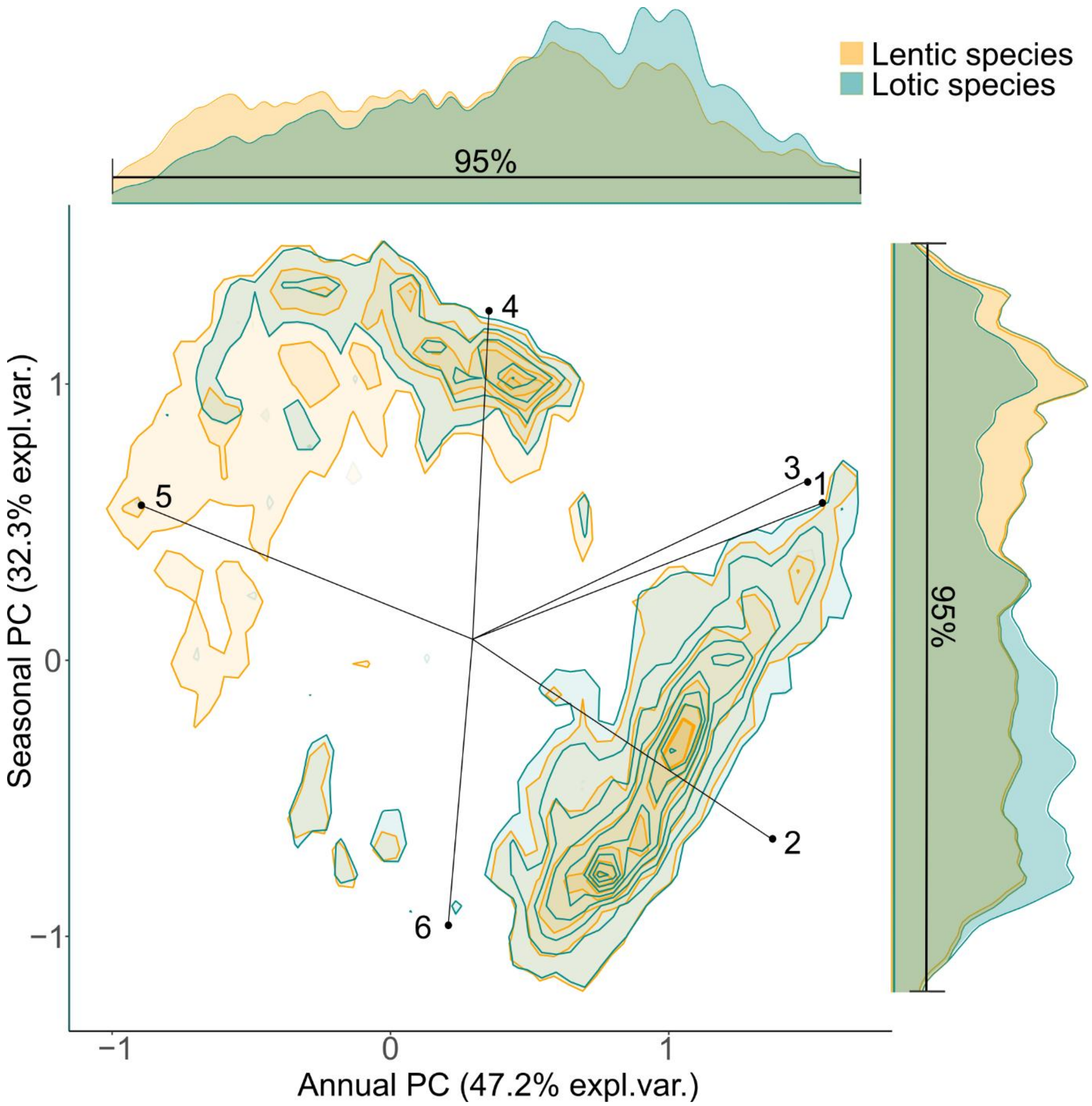


**FIGURE S3 Clustering of lentic and lotic species in the space spanned by annual and seasonal climatic drivers.** Kernel density envelopes (CI = 95%) based on species-environment combinations (*n* = 2,932) of grid cells are blue for lotic and orange for lotic species. Lotic species mainly cluster at the upper end of annual temperature and productivity gradient and the lower end of seasonality gradient, whereas lentic species occupy a greater range of both the annual and seasonal gradients. Variable loadings of Annual (first) PC and seasonal (second) principal component, respectively; (1) Annual Prod: 0.88 and 0.31; (2) Annual Temp.: 0.75 and –0.57; (3) Annual Prec.: 0.84 and 0.36; (4) Seasonality in Prod.: 0.01 and 0.65; (5) Seasonality in Temp.: –0.09 and 0.30; (6) Seasonality in Prec.: –0.09 and –0.80.

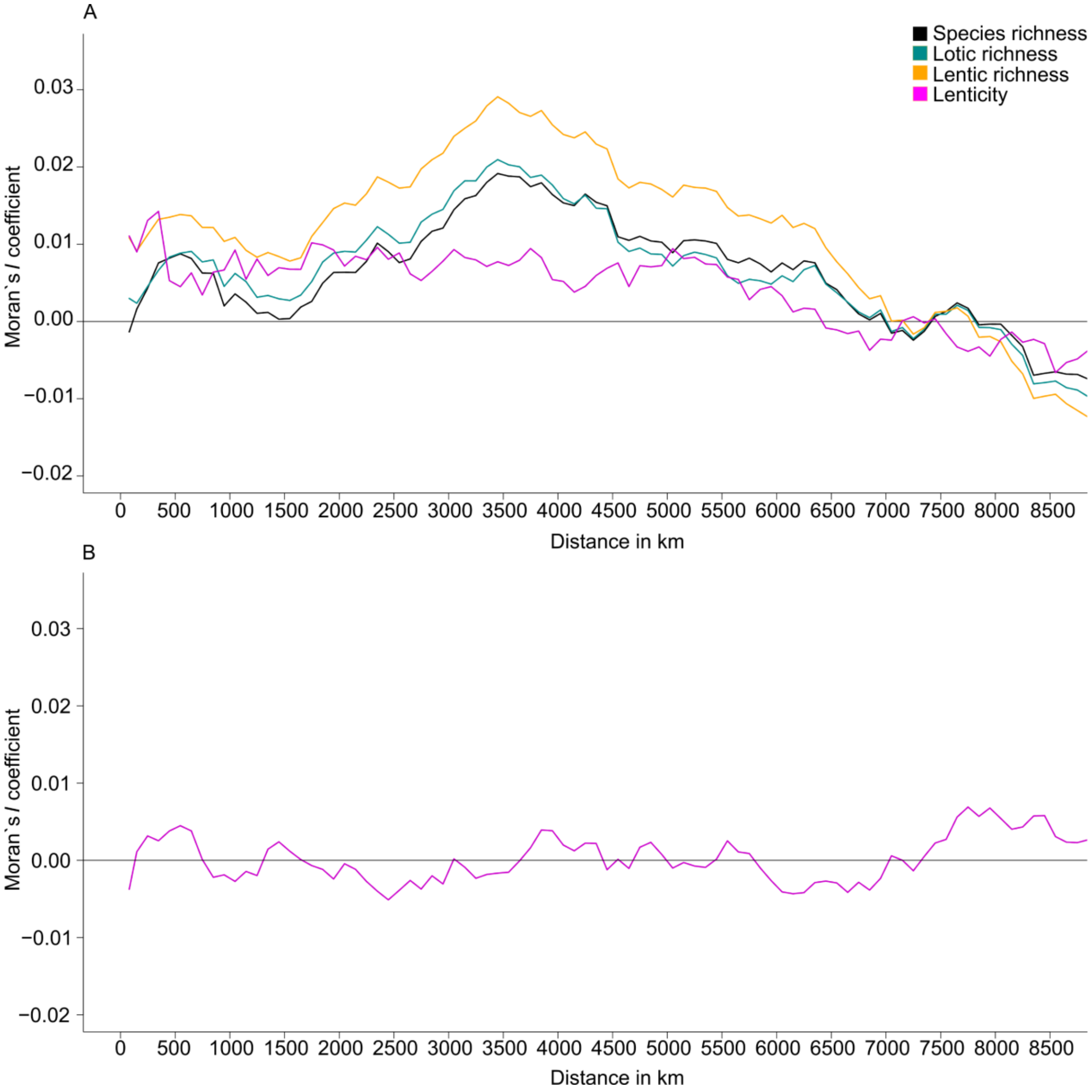


**FIGURE S4 Correlograms of residuals from ordinary least-squares regressions indicating the spatial autocorrelation (Moran's *I*).** Residuals are from regressions of observed species richness, the lentic species richness, lotic species richness (A), as well as the the proportion of lentic species (lenticity, B) with four environmental predictors each (annual PC, seasonal PC, elevation and the paleoclimatic stability, see Table 1), respectively from the regression of the proportion of lentic species with the four environmental predictors additionally with the logarithmized mean body size and range size.. The distance at which Moran's *I* reaches zero is used as the maximum threshold for the distance weight matrix in the spatial autoregressive models to account for spatial autocorrelation (see model results in Table S2).

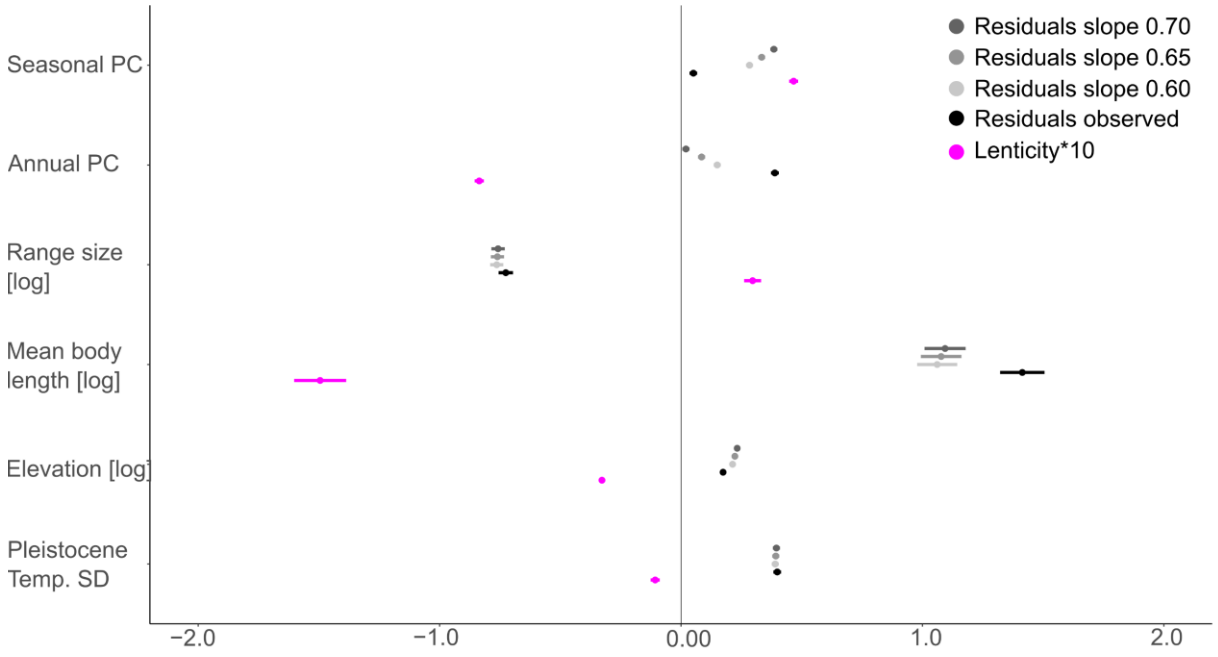


**FIGURE S5 Drivers of lenticity and residuals of species richness-temperature relationships.** Standardized effects of environmental predictors, body length and range size (coefficients and standard error bars) on the proportion of lentic species as well as residuals from models of log-transformed overall species richness and inverse mean annual temperature of odonate assemblages. Results in gray are for models fitted with the ideal slopes as predicted by the Metabolic Theory of Ecology (MTE). Results in black are effects on the residuals of the observed (MTE-naive, slope of –0.41) relationship between species richness and inverse temperature. Effects on residuals from regressions with idealized MTE slopes (i.e. –0.60, –0.65, –0.70) are gray. Lenticity was divided by 10 for visualization purposes only. Estimates of the variance explained by all six predictors or residuals of species richness-temperature relaionship and lenticity are indicated at the respective dependent variable in the legend.

#### SUPPLEMENTARY REFERENCES


1. Tarboton, W., & Tarboton, M. (2019). A Guide to Dragonflies and Damselflies of South Africa: Covering the 164 species of dragonfly and damselfly found in South Africa, Lesotho and Swaziland, 2nd ed. (Struik Nature).
2. Samways, M.J. (2008). The Dragonflies and Damselflies of South Africa (Pensoft).
3. Kalkman, V. & Orr, A. (2013). Field Guide to the Damselflies of New Guinea (NVL).
4. Orr, A.G., & Kalkman, V.J. (2015). Field Guide to the dragonflies of New Guinea (NVL).
5. Orr, A. (2005). Dragonflies of Peninsular Malaysia and Singapore (Natural History publications).
6. de Fonseka, T. (2000). The Dragonflies of Sri Lanka (Wildlife Heritage Trust Publications).
7. Haomiao, Z. (2018). Dragonflies and Damselflies of China (2-Volume Set) (Chongqing University Press).
8. Onishko, V.V. & Kosterin, O.E. (2021). Dragonflies of Russia: Illustrated Photo Guide (Fiton XXI).
9. Cho, S. (2021). Korean Odonata Adult and Larva (Gwangil Publishing Co.).
10. Marinov, M. & Ashbee, M. (2020). Dragonflies & Damselflies of New Zealand (Auckland University Press).
11. Abbott, J.C. (2011). Damselflies of Texas: A Field Guide (Illustrated Edition) (University of Texas Press).
12. Paulson, D.R. & Haber, W.A. (2021). Dragonflies and Damselflies of Costa Rica: A Field Guide (Comstock Publishing).
13. Dijkstra, K.-D.B. & Lewington, R. (2006). Field Guide to the Dragonflies of Britain and Europe (British Wildlife Publishing).
14. Paulson, D. (2011). Dragonflies and Damselflies of the East (Princeton University Press).
15. Paulson, D. (2009). Dragonflies and Damselflies of the West (Princeton University Press).
16. Bota-Sierra, C.A., Sandoval-H, J., Ayala-Sánchez, D., & Novelo-Gutiérrez, R. (2019). Dragonflies of the Colombian Cordillera Occidental, A Look from the Tatama (University of Caldas Press).
17. Garrison, R.W., von Ellenrieder, N., & Louton, J.A. (2010). Damselfly Genera of the New World (The Johns Hopkins University Press).
18. Askew, R.R. (2004). The dragonflies of Europe (Harley Books).
19. Boudot, J.-P., Borisov, S., De Knijf, G., van Grunsven, R.H.A., Schröter, A., & Kalkman, V.J. (2021). Atlas of the dragonflies and damselflies of West and Central Asia (Brill).
20. Theischinger, G., & Hawking, J. (2006). The Complete Field Guide to Dragonflies of Australia (CSIRO Publishing).
21. Kompier, T. (2015). A Guide to the Dragonflies & Damselflies of the Serra dos Órgãos (Brill).
